# Anterior Cingulate Metabolite Levels, Memory, and Inhibitory Control in Abstinent Men and Women with Alcohol Use Disorder

**DOI:** 10.1101/2022.01.07.475448

**Authors:** Emily N. Oot, Kayle S. Sawyer, Marlene Oscar-Berman, Riya B. Luhar, J.E. Jensen, Marisa M. Silveri

## Abstract

**Aims:** Alcohol use disorder (AUD), has been shown to have harmful cognitive and physiological effects, including altered brain chemistry. Further, although men and women may differ in vulnerability to the neurobiological effects of AUD, results of existing studies have been conflicting. Brain metabolite levels and cognitive functions were examined in a cross section of men with AUD (AUDm) and women with AUD (AUDw) to determine degree of abnormalities after extended periods of abstinence (mean, six years), and to evaluate gender differences in cognitive and metabolite measures.

**Methods:** Participants were 40 abstinent individuals with AUD (22 AUDw, 18 AUDm) and 50 age-equivalent non-AUD comparison participants (26 NCw, 24 NCm). Proton magnetic resonance spectroscopy (MRS) was employed at 3 Tesla to acquire metabolite spectra from the dorsal anterior cingulate cortex (dACC). Brain metabolites N-acetylaspartate (NAA), choline (Cho), myo-Inositol (mI), and glutamate & glutamine (Glx) were examined relative to measures of memory and inhibitory control.

**Results:** Metabolite levels in the AUD group showed no significant differences from the NC group. Memory and inhibitory-control impairments were observed in the AUD group. There also were significant group-specific associations between metabolite ratios and measures of inhibitory control. There were no Group-by-Gender interactions for the four metabolite ratios.

**Conclusions:** These findings demonstrate that brain metabolite levels in men and women with AUD, following long-term abstinence, do not differ from individuals without AUD. The data also provide evidence of associations between metabolite levels and measures of inhibitory control, a functional domain important for curtailing harmful drinking.

## INTRODUCTION

According to the 2019 National Survey on Drug Use and Health (NSDUH, 2020), 14.5 million people in the United States met criteria for alcohol use disorder (AUD) in the prior year. AUD is characterized by an impaired ability to control drinking despite adverse personal, societal, or occupational consequences and has been shown to have long-lasting harmful physiological and neuropsychological effects, including associations with brain abnormalities. Cognitive abilities such as general intelligence and over-learned knowledge are preserved with AUD (Stavro et al., 2013), while impairments in other functions continue despite abstinence (Jia et al., 2021, Oscar-Berman et al., 2014). Among the most common and persistent cognitive domains of impairment are memory and inhibitory control (Mullins-Sweatt et al., 2019, Stephan et al., 2017). Structural and functional magnetic resonance imaging (MRI) studies have sought to offer biological explanations linking heavy alcohol consumption and AUD with brain abnormalities as measured with magnetic resonance neuroimaging (MRI) scans (Oscar-Berman and Marinkovic, 2007, Sullivan and Pfefferbaum, 2005, Zahr et al., 2016). Likewise, AUD-related alterations in brain chemistry, measured using proton (^1^H) magnetic resonance spectroscopy (MRS) have been used to detect and quantify stability of metabolites that have important physiological functions for maintaining brain health and function (Meyerhoff et al., 2013).

The metabolite alterations widely reported in current alcohol drinkers and recently detoxified chronic heavy drinkers (Fritz et al., 2019, Meyerhoff et al., 2013) have focused on prominent, readily detectable, and reliably quantifiable brain metabolites: N-acetyl aspartate (NAA), a marker of neuronal integrity; choline (Cho), white matter integrity, membrane turnover, and inflammation; creatine (Cr), energy metabolism; myo-Inositol (mI), phospholipid metabolism and osmotic equilibrium; glutamine and glutamate (combined as Glx), neuronal activation and glucose metabolism (Govindaraju et al., 2000, Moffett et al., 2007).

Despite promising evidence of neuropsychological improvements (Oscar-Berman et al., 2014) and the associated recovery of metabolite levels after short-term abstinence (Bendszus et al., 2001), it remains unclear whether these associations are maintained after longer lengths of abstinence. Therefore, a primary objective of the current study was to employ MRS to examine NAA, mI, Cho, and Glx metabolites relative to measures of inhibitory control and memory in individuals with AUD who reported longer lengths of abstinence (average, ~6 years) than had previously been examined. It also is critical to consider possible differences in metabolite levels between men with AUD (AUDm) and women with AUD (AUDw). Although mounting evidence demonstrates an important role of gender in differentiating the impact of AUD on the brain (Nixon et al., 2014, Oscar-Berman et al., 2021, Sawyer et al., 2018), investigations of brain metabolite levels have largely been limited to men or have included insufficient numbers of women to detect group differences (Verplaetse et al., 2021). Thus, gender differences were directly examined by comparing AUDm and AUDw on metabolite levels and on their associations with length of abstinence and neuropsychological measures of memory and inhibitory control.

## METHODS

### Participant recruitment and inclusion

The study included 40 abstinent individuals with AUD (22 AUDw, 18 AUDm) and 50 non-AUD age-matched comparison subjects (26 NCw, 24 NCm). Participants were recruited from the Boston area via newspaper and web-based advertisements and flyers. The study was approved by the participating Institutional Review Boards: Boston University School of Medicine (#H24686), VA Boston Healthcare System (#1017 and #1018), and Massachusetts General Hospital (#2000P001891). Participants provided written informed consent and were reimbursed $15 per hour for assessments, $25 per hour for scans, and $5 for travel expenses.

Participants were interviewed to ensure they met inclusion criteria. Participants were given the Computerized Diagnostic Interview Schedule (Robins et al., 2000), which provides psychiatric diagnoses according to criteria established by the American Psychiatric Association (DSM-IV; (APA, 1994)). Participants were excluded if English was not one of their first languages, if they were predominantly left-handed, or if they had any of the following: Korsakoff’s syndrome; HIV; cirrhosis; major head injury with loss of consciousness greater than 15 minutes unrelated to AUD; stroke; epilepsy or seizures unrelated to AUD; history of electroconvulsive or electro-shock therapy; major neurotoxicant exposure; psychotic disorders; bipolar II; Hamilton Rating Scale for Depression (HRSD (Hamilton, 1960)) score over 18; or history of drug use once per week or more within the past five years, except for two cases: one AUD woman who had used marijuana regularly (but not within the past three months), and one NC man who had used marijuana regularly (but not within the past two years). The AUD participants were excluded if they did not have positive criteria for alcohol abuse or dependence, if their duration of heavy drinking was less than five years, or if they had consumed alcohol within four weeks prior to testing. Non-AUD control participants were excluded if they reported duration of heavy drinking greater than one year. One participant was excluded for claustrophobia; another was excluded after a brain lesion was identified; and 23 were excluded for unusable LCModel fits of spectroscopy data.

### Participant assessment procedures

Neuropsychological measures of the domains of memory and inhibitory control were examined. Memory was assessed using the Working Memory subtest of the Wechsler Adult Intelligence Scale - Fourth Edition (WAIS-IV) and the Immediate Memory and Delayed Memory subtests of the Wechsler Memory Scale - Fourth Edition (WMS-IV) (Holdnack and Drozdick, 2010). Inhibitory control was assessed using Dickman’s Impulsivity Inventory (Dickman, 1990) and the Barratt Impulsiveness Scale (BIS-11, (Patton et al., 1995)). Three Delis-Kaplan Executive Function System measures (D-KEFS; (Delis et al., 2001)) were acquired: Trails Number Sequencing-2 (analogous to Trails A), Trails Number-Letter Switching-4 (analogous to Trails B) and Verbal Letter Fluency-1 (analogous to Controlled Oral Word Association Test (COWAT)) (Lezak et al., 2004). For the D-KEFS, scaled scores were used, and higher scores indicate better performance.

Participants were administered the Alcohol Use Questionnaire (Cahalan et al., 1969), which yields length of sobriety (LOS; in years), duration of heavy drinking (DHD; in years), and a quantity frequency index (ounces of alcohol per day, roughly equivalent to daily drinks [DD]). The LOS refers to the period between the MRI scan date and the last reported drink. The DHD represents the total number of years participants drank at least 21 drinks per week (average three drinkers per day). The DD measure reflects the last six months (NC group), or during the six months preceding cessation of drinking (AUD group). Five AUD participants drank fewer than three drinks/day during the six months prior to cessation; thus, DD was obtained for the last six months of heavy drinking. All participants also completed the Alcohol Use Disorders Identification Test (AUDIT) (Babor et al., 1992). The questionnaire was modified to be past tense for the AUD group, to assess time during which they were drinking heavily.

### MRI acquisition

MRI scans were obtained at Massachusetts General Hospital on a 3 Tesla Siemens MAGNETOM Trio Tim scanner with a 32-channel head coil (123.18MHz). Once positioned, head placement was confirmed using three-plane scout images. Two T1 weighted multi-echo MP-RAGE series for volumetric analysis (one AUD man had only one series) were obtained with these parameters: TR=2530ms, TE=1.79ms, 3.71ms, 5.63ms, 7.55ms (RMS average used), flip angle=7°, field of view=256mm, matrix=256×256, slice thickness=1mm, 176 interleaved sagittal slices, with GRAPPA acceleration factor of 2.

### Magnetic Resonance Spectroscopy

A 20mm x 20mm x 20mm oblique voxel was prescribed in the anterior cingulate cortex (ACC) along the midline, with the inferior plane of the voxel parallel to the descending surface of the corpus callosum (Figure 1). Voxel shimming, flip-angle, water-suppression and frequency were automatically adjusted using Siemens software. Proton spectroscopy data were acquired using a Point-Resolved Echo Spectroscopy Sequence (PRESS) to acquire water-suppressed TE=30ms ACC spectra. Additional acquisition parameters included TR=2s, spectral bandwidth=1.2kHz, readout duration=512ms, NEX=128, total scan duration=4.3min. Spectroscopic data processing was performed using in-house reconstruction code and LCModel (Provencher, 1993). The 30ms LCModel basis set utilized a GAMMA-simulated model based on the PRESS sequence. Five metabolite levels were assessed: Cho-containing compounds, Glx, mI, NAA, and total Cr (tCr, Cr + phosphocreatine). Calculating metabolite ratios relative to tCr as an internal standard, a method that has inherent limitations, is accepted in the field as a validated and reliable method for examining metabolites (Jansen et al., 2006). Thus, ratios of metabolites to tCr were calculated: Cho/tCr, Glx/tCr, mI/tCr, and NAA/tCr.

**Figure 1.**
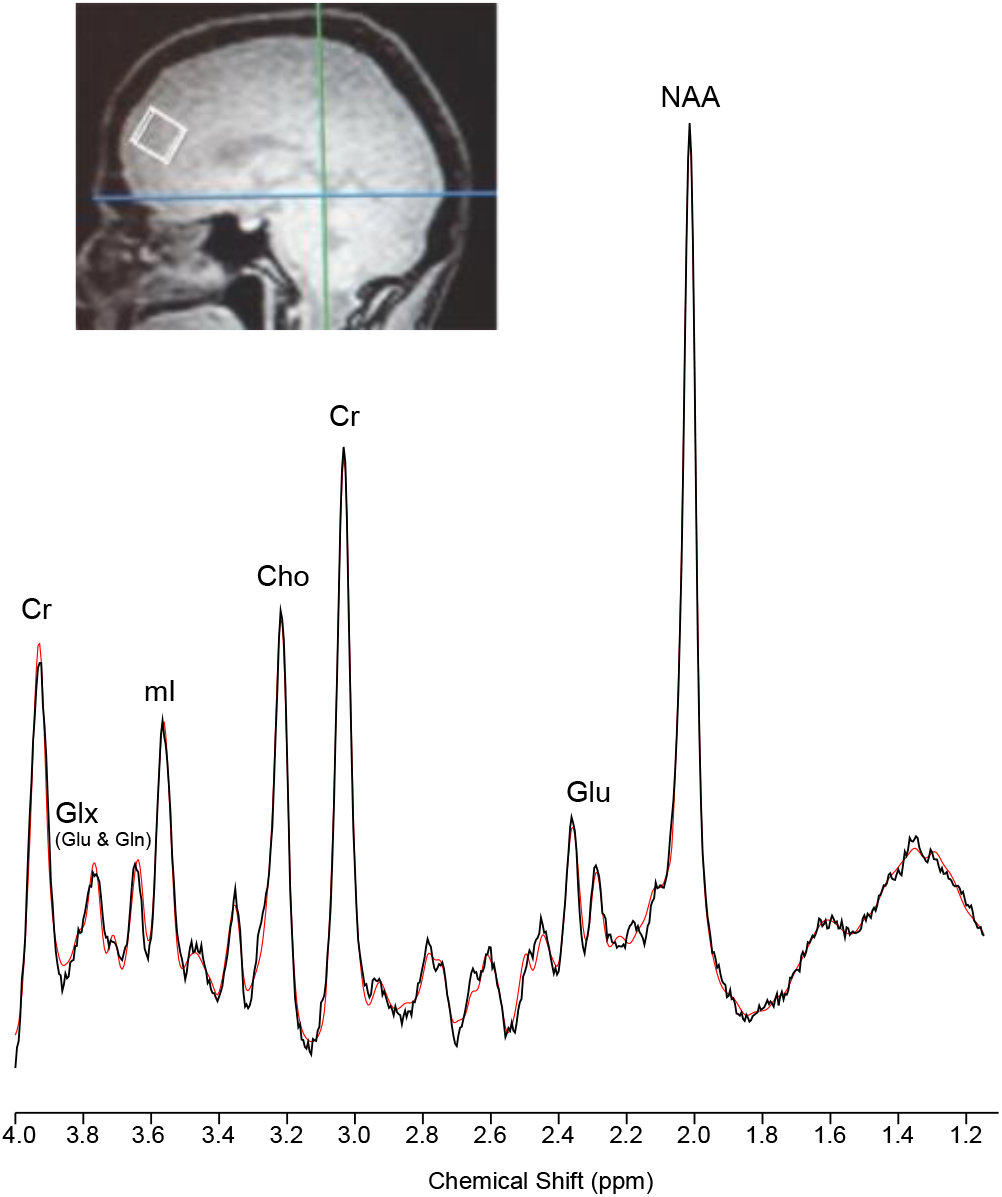
MRS Voxel Placement and Sample Proton Spectrum. Sagittal image illustrating the placement of 20mm x 20mm x 20 mm single voxel in the ACC and a sample spectrum (black) and LCModel fit (red) from a study participant. Abbreviations: ACC, anterior cingulate cortex; tCr, creatine; Cho, choline; Glu, glutamate; Glx, glutamate and glutamine; NAA, N-acetyl-aspartate; mI, myo-Inositol; ppm, parts per million.

### Image segmentation analysis

The high-resolution image sets (T1-weighted) were segmented into gray matter (GM), white matter (WM), and cerebrospinal fluid (CSF) binary-tissue maps (FSL, Oxford, UK). Partial tissue percentages were extracted for the oblique ACC voxel for use in analyses.

### Statistical analyses

Statistical analyses were performed using R version 3.4.0 (R Core Team, 2020). To examine Group-by-Gender interactions, separate general linear models were constructed using the lm function in R, predicting each measure of interest: the participant characteristics (Table 1), and the tissue contributions and four metabolite ratios (Table 2 and Figure S1). Regression assumptions were visually confirmed (normality of outcome measures, homogeneity of variance), and thresholds were set for multicollinearity (Pearson correlations < 0.5) and influence (Cook’s D < 1.0). For each model, findings are reported from the ANCOVA (using the car:Anova function with Type III sums of squares), followed by the results from the post hoc analyses (using the emmeans package; (Lenth, 2016)). The interaction of Group-by-Gender was examined to assess how the impact of AUD differed for men and women.

**Table 1.**
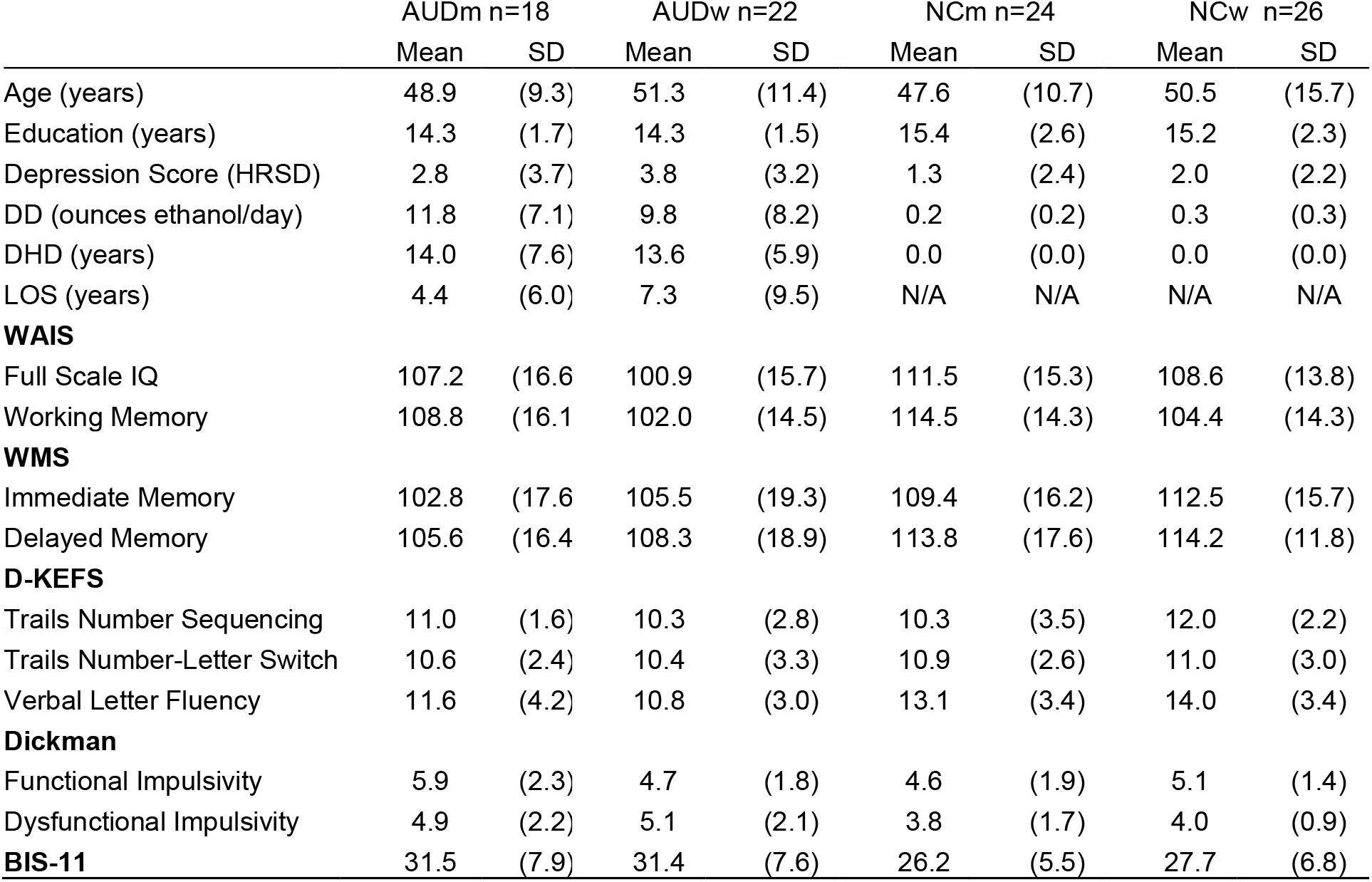
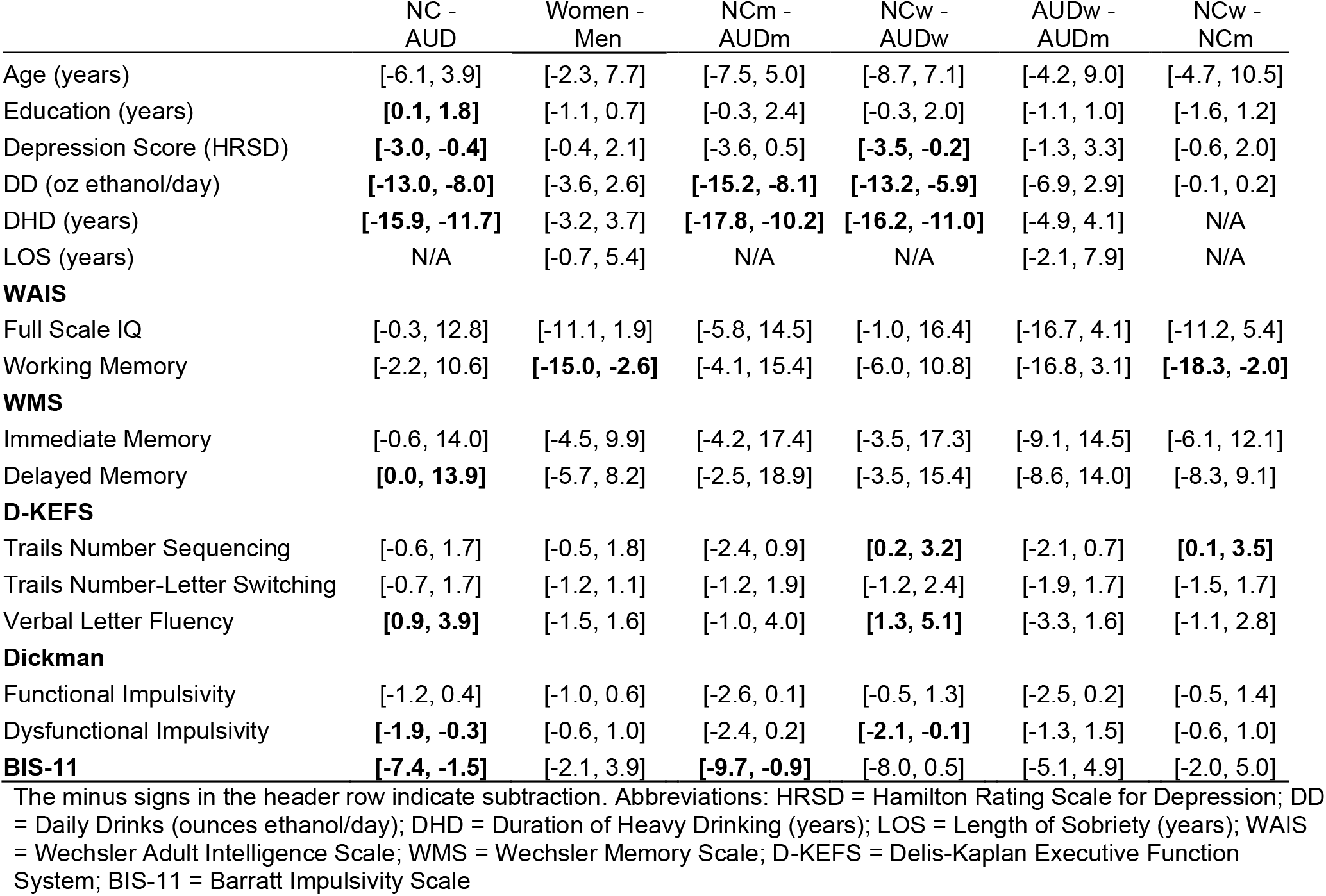
Participant characteristics.

**Table 2.** Metabolite ratios (normalized to creatine levels), and tissue contributions.

|  | AUDm<br>n=18 |  | AUDw<br>n=22 |  | NCm<br>n=24 |  | NCw<br>n=26 |  |
| --- | --- | --- | --- | --- | --- | --- | --- | --- |
|  | Mean | SD | Mean | SD | Mean | SD | Mean | SD |
| Cho/tCr | 0.25 | 0.03 | 0.23 | 0.03 | 0.25 | 0.03 | 0.24 | 0.03 |
| Glx/tCr | 0.98 | 0.14 | 1.05 | 0.12 | 1.02 | 0.15 | 1.04 | 0.14 |
| ml/tCr | 0.80 | 0.12 | 0.76 | 0.11 | 0.78 | 0.10 | 0.77 | 0.10 |
| NAA/tCr | 1.07 | 0.17 | 1.08 | 0.14 | 1.09 | 0.18 | 1.09 | 0.16 |
| GM | 57.77 | 6.29 | 59.27 | 6.84 | 59.19 | 7.55 | 59.49 | 8.49 |
| WM | 26.65 | 4.92 | 26.24 | 5.61 | 26.56 | 5.87 | 27.34 | 7.11 |
| CSF | 15.81 | 4.62 | 15.24 | 5.59 | 13.01 | 5.16 | 12.60 | 4.18 |

|  | NC - AUD<br>95% CI | Women - Men<br>95% CI | NCm - AUDm<br>95% CI | NCw - AUDw<br>95% CI | AUDw - AUDm<br>95% CI | NCw - NCm<br>95% CI |
| --- | --- | --- | --- | --- | --- | --- |
| Cho/tCr | [-0.01, 0.01] | **[-0.02, 0.00]** | [-0.02, 0.02] | [-0.02, 0.02] | [-0.03, 0.00] | [-0.03, 0.00] |
| Glx/tCr | [-0.05, 0.06] | [-0.02, 0.10] | [-0.06, 0.13] | [-0.09, 0.06] | [-0.02, 0.15] | [-0.06, 0.10] |
| ml/tCr | [-0.05, 0.04] | [-0.07, 0.02] | [-0.09, 0.05] | [-0.05, 0.07] | [-0.11, 0.03] | [-0.07, 0.05] |
| NAA/tCr | [-0.05, 0.08] | [-0.06, 0.07] | [-0.10, 0.12] | [-0.07, 0.10] | [-0.10, 0.10] | [-0.09, 0.10] |
| GM | [-2.29, 3.80] | [-2.27, 3.88] | [-2.91, 5.74] | [-4.23, 4.68] | [-2.72, 5.70] | [-4.26, 4.86] |
| WM | [-1.92, 3.00] | [-2.25, 2.72] | [-3.46, 3.28] | [-2.59, 4.80] | [-3.79, 2.96] | [-2.92, 4.48] |
| CSF | **[-4.77, -0.63]** | [-2.52, 1.72] | [-5.86, 0.27] | [-5.57, 0.28] | [-3.83, 2.70] | [-3.11, 2.28] |

The minus signs in the header row indicate subtraction. The upper set of numbers provides means and standard deviations, and the lower numbers represent 95% confidence intervals (CI). Abbreviations: NAA = N-acetylaspartate; Cho = choline-containing compounds; ml = myo-Inositol; Glx = glutamate + glutamine; tCr = total creatine; GM = gray matter; WM = white matter; CSF = cerebrospinal fluid. Bolded numbers indicate significance at $p < 0.05$ .

Five categories of regression analyses were conducted, as detailed below. For Group and Gender differences (1) participant characteristics, (2) GM, WM, and CSF tissue contributions, and (3) metabolite ratio levels were examined. Relationships between metabolite levels, (4) drinking history, and (5) neuropsychological measures also were examined.

1. Group and Gender differences for participant characteristics were assessed with a model that included the interaction of Group and Gender (for example, specified as “age ~ group*gender”).
2. Group and Gender differences for tissue contributions were assessed with a model that included the interaction of Group and Gender.
3. Group and Gender differences for metabolite ratios were assessed with a model that included the interaction of Group and Gender, with GM and education as covariates (for example, specified as “choline_ratio ~ group*gender + GM + education”). Age was highly correlated with GM (r = 0.5), so it was not included as a covariate.
4. Relationships of metabolite levels to the three measures of drinking history (DHD, DD, and LOS) were assessed in the AUD group. For each of the four metabolites, a model was constructed to assess effects of drinking history measures and interactions with Gender (for example, specified as “choline_ratio ~ DHD*gender + DD*gender + LOS*gender + GM + education”).
5. Relationships between metabolites and neuropsychological measures were assessed. For each of the four metabolites, separate models were constructed for each of the 10 measures: WAIS-IV Full Scale IQ, WAIS-IV Working Memory, WMS-IV Immediate Memory, WMS-IV Delayed Memory, D-KEFS Trails Number Sequencing-2, D-KEFS Trails Number-Letter Switching-4, D-KEFS Verbal Letter Fluency-1, Dickman Functional Impulsivity, Dickman Dysfunctional Impulsivity, and BIS-11. Each model was specified to examine interactions of the neuropsychological measure with Group and Gender while accounting for gray matter tissue contribution (for example, “choline_ratio ~ bis*alcoholism*gender + GM”), for a total of 40 models. Age and education were not included as covariates because normative scaled scores were used. For all analyses, significant predictors and interactions indicated by ANCOVAs were followed by post-hoc comparisons using the emmeans::emtrends function. This function allows slopes within the model to be compared (for example, how relationships of NAA/tCr with BIS differ for the AUD vs. NC groups).

In addition to regressions, correlational analyses were conducted to examine how drinking history related to memory and inhibitory control measures in the AUD group, using the ppcor::pcor.test function to calculate partial correlation values for relationships between metabolite ratios and neuropsychological measures for each group separately, accounting for GM tissue contribution.

## RESULTS

### Participant characteristics

Participant characteristics are summarized in Table 1. The AUD group had a mean LOS of 6.0 years (range 0.1 to 32.8), mean DD of 10.7 drinks (range 2.3 to 34.8), and mean DHD of 13.8 drinks (5.0 to 32.0). Analyses of Group (AUD vs. NC) and Gender revealed no significant main effects or interactions with age (NC mean=49.1 years, range=26.6 to 77.0 years; AUD mean=50.2 years, range=28.2 to 73.3 years). There was a significant main effect of Group for education (*F*_1,86_=4.40, *p*=0.04), with education of the NC group (mean=15.3 years) being higher than the AUD group (mean=14.3 years), but there was no significant main effect of Gender or Group-by-Gender interaction. HRSD scores were significantly higher, 0.4 to 3.0 points (95% CI) in the AUD group (mean=3.3) compared to the NC group (mean=1.6) (*F*_1,86_=7.56, *p*=0.007). Analyses of Gender effects for drinking history variables in the AUD group revealed no significant main effects of Gender.

Analyses of memory measures demonstrated that women had lower scores on the WAIS Working Memory Scale than men (*F*_1,86_=7.36, *p*=0.008) and that the AUD group had lower scores on the WMS Delayed Memory than the NC group (*F*_1,86_=4.21, *p*=0.04) (Table 1). Analyses of inhibitory control measures showed a Group-by-Gender interaction for D-KEFS Number Sequencing-2 (*F*_1,86_=4.75, *p*=0.03), with NCw exhibiting higher scores than AUDw, while NCm scored significantly lower than AUDm. The AUD group had lower D-KEFS Verbal Letter Fluency-1 scores than the NC group (*F*_1,86_=10.06, *p*=0.002). There was a significant Group-by-Gender interaction for Functional Impulsivity (*F*_1,86_=4.56, *p*=0.04); the AUDm had a higher score than NCm, but AUDw had significantly lower scores than NCw. The AUD group also demonstrated significantly higher scores than the NC group on the Dickman Dysfunctional Impulsivity Scale (*F*_1,86_=8.59, *p*=0.004) and on the BIS-11 (*F*_1,86_=9.33, *p*=0.003).

Partial correlational analyses of memory and inhibitory control measures relative to drinking history in the AUD group demonstrated a significant positive correlation between DHD and Dickman Dysfunctional Impulsivity scores (*r*_40_=0.40, *p*=0.01). Additionally, LOS in the AUD group was positively related to Age (*r*_40_=0.50, *p*=0.001) and Education (*r*_40_=0.33, *p*=0.04), and negatively related to DD (*r*_40_=-0.31, *p*=0.05). LOS also was positively associated with WAIS Working Memory (*r*_40_=0.33, *p*=0.04) and Immediate Memory (*r*_40_=0.39, *p*=0.01).

### Metabolite ratios and tissue contributions

Analyses of Group and Gender effects for tissue contributions revealed no significant main effects nor interactions for GM or WM. However, CSF levels were higher in the AUD group relative to the NC group (Table 2).

Metabolites for the AUD and NC groups were not statistically different. The main effect of Gender for the Cho/tCr ratio was significant (*F*_1,84_=5.04, *p*=0.03), with higher Cho/tCr in men than in women. No other significant main effects of Group or Gender, or Group-by-Gender interactions were observed (Table 2).

### Associations between metabolite ratios and neuropsychological measures

Regression models were constructed to identify how relationships between metabolite ratios and neuropsychological scores differed by Group. There were four associations with slopes that were significant within a group and significantly different from the corresponding (non-significant) relationship in the other group (Table 3 and Figure 2). The AUD group had stronger relationships (steeper slopes) than the NC group for three measures, as follows. (1) For Dickman Functional Impulsivity, the AUD group had a significantly more positive slope in relation to Cho/tCr (*t*_81_=2.2, *p*=0.03). (2) For Dickman Dysfunctional Impulsivity, the AUD group had a significantly more negative slope in relation to mI/tCr (*t*_81_=-2.4, *p*=0.02). (3) For the BIS-11, the AUD group had a significantly more positive slope in relation to NAA/tCr than the NC group (*t*_81_=2.9, *p*=0.005). The NC group demonstrated stronger relationships than the AUD group for D-KEFS Trails Number Sequencing, with Glx/tCr (*t*_81_=-2.0, *p*=0.05). Additionally, compared to women, men had a stronger association between scores on the Dickman Functional Impulsivity and mI/tCr ratios (*t*_81_=2.3, *p*=0.03).

**Figure 2.**
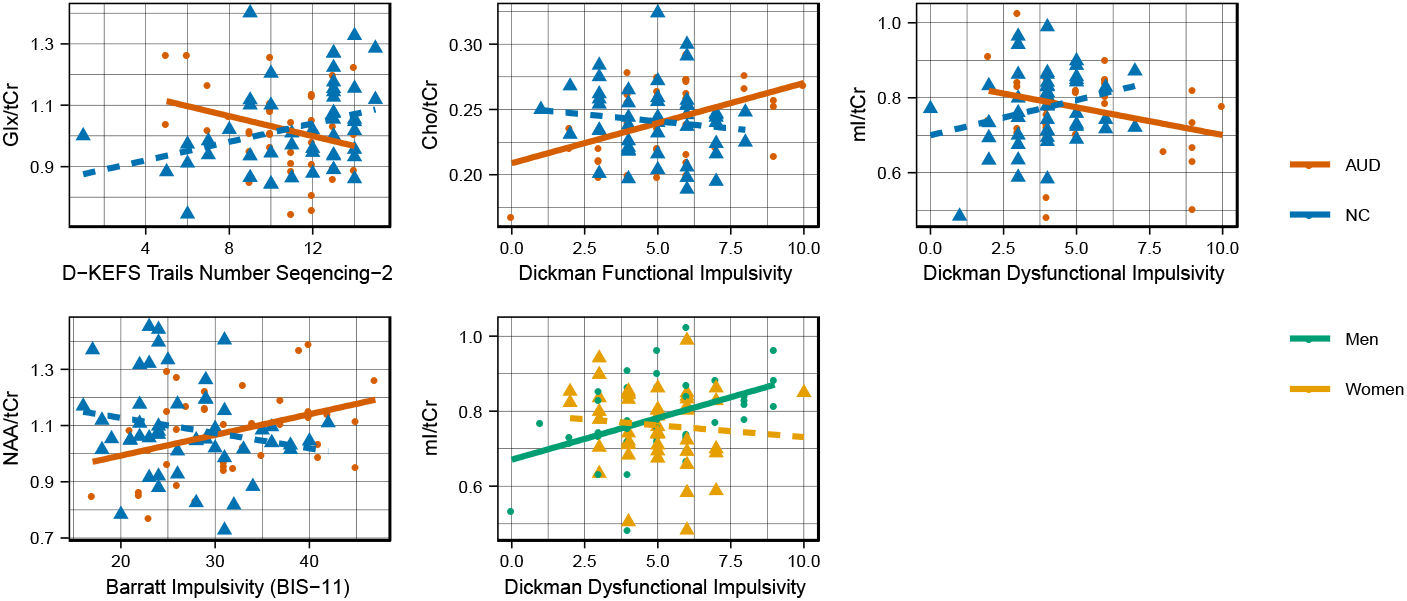
Relationships between metabolite ratios and neuropsychological measures. Abbreviations: AUD = Alcohol Use Disorder; NC = Non-AUD Control; NAA = N-acetyl-aspartate; Cho = choline-containing compounds; mI = myo-Inositol; Glx = glutamate + glutamine; tCr = total creatine

**Table 3.** Metabolite ratios: Interactions of group with neuropsychological measures.

| Metabolite Ratio | Measures of Inhibitory Control | Group Interaction<br>F(1, 81) | AUD<br>95% CI | r | NC<br>95% CI | r |
| --- | --- | --- | --- | --- | --- | --- |
| Glx/tCr | D-KEFS Trails Number Sequencing-2 | **6.9** | [-0.03, 0.01] | -0.3 | **[0.00, 0.03]** | **0.3** |
| Cho/tCr | Dickman Functional Impulsivity | **4.2** | **[0.00, 0.01]** | **0.4** | [-0.01, 0.00] | -0.1 |
| ml/tCr | Dickman Dysfunctional Impulsivity | **6.5** | **[-0.03, 0.00]** | **-0.3** | [-0.01, 0.04] | 0.2 |
| NAA/tCr | BIS-11 | **7.0** | **[0.00, 0.01]** | **0.4** | [-0.01, 0.00] | -0.2 |

| Metabolite Ratio | Measures of Inhibitory Control | Gender Interaction<br>F(1,81) | Men<br>95% CI | r | Women<br>95% CI | r |
| --- | --- | --- | --- | --- | --- | --- |
| ml/tCr | Dickman Functional Impulsivity | **5.1** | **[0.01, 0.04]** | **0.4** | [-0.03, 0.01] | -0.1 |

## DISCUSSION

The main results of this project are threefold. First, by demonstrating that proton metabolites measured in the ACC of long-term abstinent men and women with AUD are similar to those of individuals without AUD, the current results extend findings from prior studies that demonstrated recovery of metabolite levels after shorter term periods of detoxification and abstinence. Second, significant associations between metabolite levels and measures of inhibitory control in individuals with AUD were observed, suggesting continued functional relevance of these metabolites in AUD even after sustained abstinence. And third, possible differences in metabolite levels between AUDm and AUDw were examined, though none were observed.

The equivalence in metabolite levels observed between AUD and NC groups (within 10%) is an encouraging sign for current or recently detoxified heavy drinkers, suggesting that alterations in metabolite levels, which are thought to be markers of acute brain injury (Meyerhoff et al., 2013), are reversible and may recover completely and permanently, even when some volumetric or functional brain differences persist. This finding is consistent with previous MRS studies documenting normalization of brain metabolites after shorter periods of abstinence. The AUD cohort examined in this study had been abstinent for a long duration (mean 6 years), thus these results contribute to the characterization of the timeline of metabolite recovery by demonstrating that improvements in metabolite levels seen in recently detoxified heavy drinkers persist with long-term sobriety.

Indeed, the AUD group had impairments in measures of both memory (WMS Delayed Memory) and inhibitory control (Dickman Dysfunctional Impulsivity, and BIS-11), despite there being no substantial group differences from the NC group in metabolite levels. This finding supports the idea that differences seen in long-term abstinent AUD individuals are more likely to stem from persistent structural brain abnormalities (Monnig et al., 2013) or from pre-existing personality factors (Mullins-Sweatt et al., 2019, Ruiz et al., 2013) than they are to stem from the acute disruption of brain chemistry associated with active alcohol consumption or withdrawal. For example, both impulsivity (Stephan et al., 2017) and poor memory may be risks for heavy drinking (Verdejo-Garcia and Bechara, 2009).

Several significant correlations were observed between measures of drinking history and the neuropsychological measures in the AUD group. There was a significant positive correlation between DHD and Dickman Dysfunctional Impulsivity scores, indicating that people with longer heavy drinking histories have more dysfunctional behavior patterns. Additionally, LOS was positively related to Age and Education, and negatively related to DD, indicating that individuals in the current sample who had maintained their sobriety longest were older (perhaps a direct function of the years required to have a longer LOS), better educated, and drank less when they were actively drinking. LOS also was positively associated with WAIS Working Memory, and with WMS Immediate Memory, suggesting recovery of neuropsychological functions with abstinence, or perhaps a protective effect of memory skills in maintaining sobriety.

These data provide preliminary evidence of associations between metabolite ratios and scores of the three impulsivity measures that are significantly more pronounced in the AUD group than in the NC group. NAA was related to higher impulsivity on the BIS-11 in the AUD group, perhaps implicating a relationship between ACC neuronal integrity and more impulsive actions for people who have a history of heavy drinking. Higher functional impulsivity in the AUD group was associated with higher Cho/tCr ratios, suggesting a relationship between spontaneous behavior patterns and ACC membrane turnover in this same population. Dysfunctional Impulsivity was associated with lower mI/tCr in the AUD group, which might suggest less ACC membrane turnover or inflammation despite more behavioral impairment in this population. While these metabolite levels do not have established relationships with neurobehavioral markers such as impulsivity, these relationships point to subtle influences of each metabolite.

For Trails Number Sequencing, the relationship to Glx/tCr was stronger in the NC group than in the AUD group. This metabolite ratio has been associated with tissue health, so it follows that lower scores on a measure of control would be associated with lower levels of tissue health in the ACC in the NC group. The poorer scores in some participants of the AUD group despite the higher ratio of Glx/tCr might point to other (unmeasured) forms of dysfunction in the region.

Finally, there have been few prior studies that have included sufficient sample sizes of AUDw to permit examination of gender differences in metabolites measured by proton spectroscopy. The identification of gender differences is a crucial component of precision medicine and will contribute to the understanding of how addiction differs between people, depending on individual factors like age and gender. Armed with that knowledge, clinicians can better tailor treatment and prevention strategies. In this study, the proportion of women (48 women of 90 participants) was substantially larger than in prior MRS studies examining metabolite levels in the context of AUD. While the results indicated similar levels of NAA/tCr, mI/tCr, and Gx/tCr in men and women, Cho/tCr was significantly higher in men (by ~6% or 0.4 SD). Cho/tCr is known to be associated with white matter integrity, no relationship was observed with LOS nor with behavioral measures, suggesting this gender difference may reflect differences in brain composition.

Two significant Group-by-Gender interactions were identified for measures of inhibitory control: Dickman functional impulsivity and Trails number sequencing. For both measures, the impact of AUD was opposite for the two genders: The NCw scored higher than AUDw, but NCm scored lower than AUDm. These findings indicate that AUDm were more impulsive but also had faster processing speed than AUDw, consistent with previous reports (Fama et al., 2019, Stoltenberg et al., 2008).

Finally, a significant gender interaction was evident for functional impulsivity predicting mI/tCr, wherein the relationship was more pronounced in men than women. Preliminary speculation might suggest that higher levels of mI/tCr in the ACC interact with elements of the brain or hormones in men to produce higher impulsivity, whereas the higher levels in women do not have that effect. Hormone levels are modulated by stress, an important factor during the progression of AUD in men and women (Peltier et al., 2019).

An important limitation of the present research is that, due to the cross-sectional nature of the study, longitudinal changes over the course of long-term abstinence was not to be determined. Nonetheless, these data are useful when interpreted in conjunction with existing longitudinal datasets of metabolite levels and drinking behaviors in shorter-term abstinence (Fritz et al., 2019, Meyerhoff et al., 2013), and provide important insight into what sustained abstinence means for ACC neurochemistry and neuropsychological outcomes. Moreover, further longitudinal studies can help to establish causal links between long-term duration of abstinence and metabolite ratios, and to elucidate relationships among metabolite levels and neuropsychological measures across groups. Another limitation is that the potential impact of tobacco use on the metabolite ratios in the current sample was not examined. Smoking can contribute to alterations seen in metabolite ratios (Durazzo et al., 2004), and can have meaningful implications for neuroimaging results (Luhar et al., 2013). Also, the use of tCr as a normalizing denominator, while a common practice, can limit the interpretability of results that use this approach (Tunc-Skarka et al., 2015, Zahr et al., 2016). Alternative normalization procedures, such as external referencing or water signal normalization, also have drawbacks that can confound results.

In conclusion, these findings demonstrate that brain metabolite levels in men and women with AUD, following long-term abstinence, do not differ from individuals without AUD. The data also provide evidence of associations between metabolite levels and measures of inhibitory control in the population of men and women with AUD in long-term abstinence. There also were no substantial differences in metabolite levels between men and women with AUD, suggesting that if metabolites are normalized to non-AUD levels, the current data do not support the notion that women have longer term impacts related to AUD on neurochemistry.

## Acknowledgments

This work was supported by funds from the National Institute on Alcohol Abuse and Alcoholism (NIAAA) grants F31AA025824 to Emily Oot, R01AA007112 and K05AA00219 to Marlene Oscar Berman, and NIAAA K24 AA025977, R01AA018153, and K01AA014651 to Marisa Silveri, and by the US Department of Veterans Affairs Clinical Science Research and Development grant I01CX000326 to Dr. Marlene Oscar Berman. The authors thank Zoe Gravitz, Gordon Harris, Steve Lehar, Pooja Parikh Mehra, Diane Merritt, Susan Mosher Ruiz, and Maria Valmas for assistance with recruitment, assessment, data analyses, neuroimaging, or manuscript preparation. We also wish to acknowledge the Athinoula A. Martinos Center of Massachusetts General Hospital for imaging resources (1S10RR023401, 1S10RR019307, and 1S10RR023043). Finally, we would like to acknowledge the role of the research participants for making this study possible. The content is solely the responsibility of the authors and does not necessarily represent the official views of the National Institutes of Health, the U.S. Department of Veterans Affairs, or the United States Government.

---

**Figure S1.**
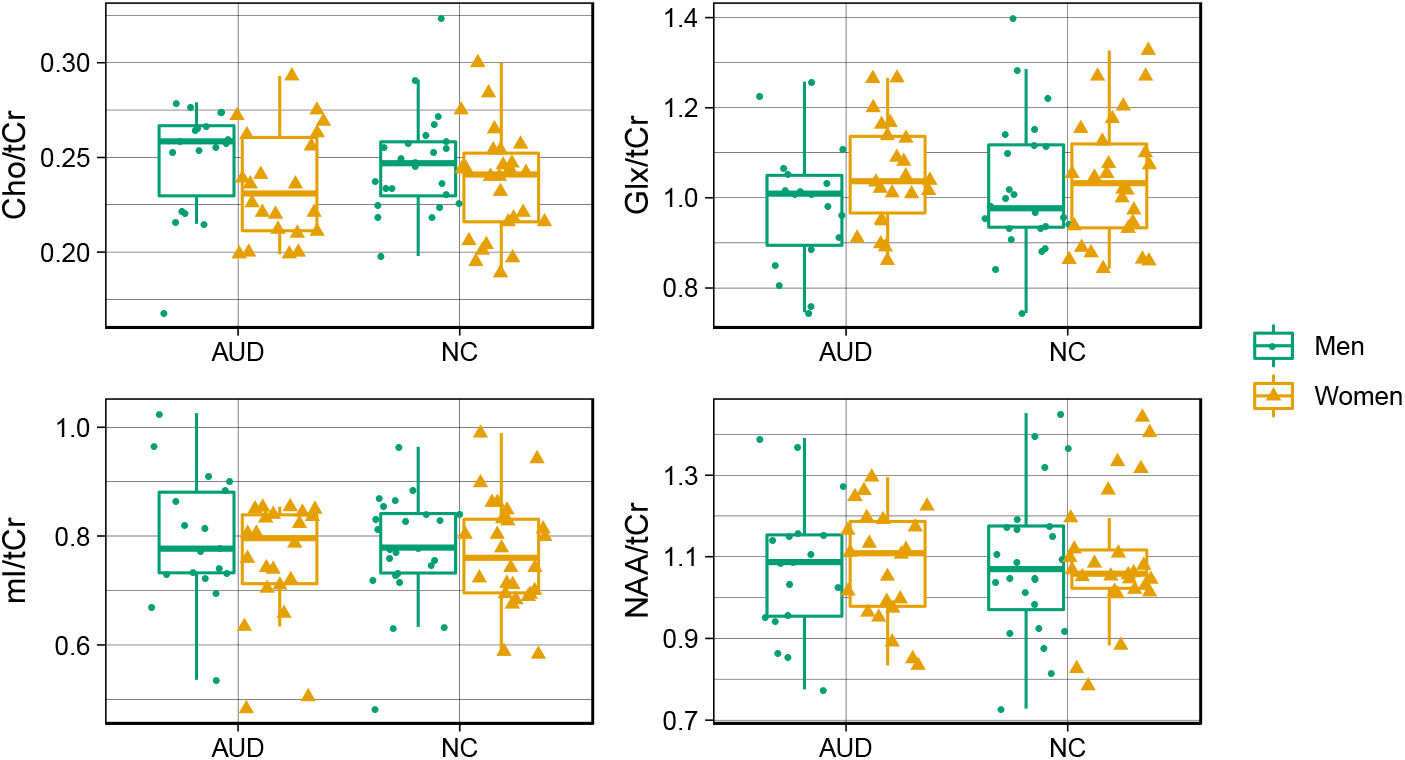
Metabolite levels. Abbreviations: AUD = Alcohol Use Disorder; NC = Non-AUD Control; NAA = N-acetyl aspartate; Cho = choline-containing compounds; mI = myo-Inositol; Glx = glutamate; tCr = total creatine.

